# Sirtuin 2 inhibition modulates chromatin landscapes genome-wide to induce senescence in ATRX-deficient malignant glioma

**DOI:** 10.1101/2023.01.09.523324

**Authors:** Prit Benny Malgulwar, Carla Danussi, Sharvari Dharmaiah, William E. Johnson, Arvind Rao, Jason T. Huse

## Abstract

Inactivating mutations in *ATRX* characterize large subgroups of malignant gliomas in adults and children. ATRX deficiency in glioma induces widespread chromatin remodeling, driving transcriptional shifts and oncogenic phenotypes. Effective strategies to therapeutically target these broad epigenomic sequelae remain undeveloped. We utilized integrated mulit-omics and the Broad Institute Connectivity Map (CMAP) to identify drug candidates that could potentially revert ATRX-deficient transcriptional changes. We then employed disease-relevant experimental models to evaluate functional phenotypes, coupling these studies with epigenomic profiling to elucidate molecular mechanim(s). CMAP analysis and transcriptional/epigenomic profiling implicated the Class III HDAC Sirtuin2 (Sirt2) as a central mediator of ATRX-deficient cellular phenotypes and a driver of unfavorable prognosis in ATRX-deficient glioma. Sirt2 inhibitors reverted Atrx-deficient transcriptional signatures in murine neuroprogenitor cells (mNPCs) and impaired cell migration in Atrx/ATRX-deficient mNPCs and human glioma stem cells (GSCs). While effects on cellular proliferation in these contexts were more modest, markers of senescence significantly increased, suggesting that Sirt2 inhibition promotes terminal differentiation in ATRX-deficient glioma. These phenotypic effects were accompanied by genome-wide shifts in enhancer-associated H3K27ac and H4K16ac marks, with the latter in particular demonstrating compelling transcriptional links to Sirt2-dependent phenotypic reversals. Motif analysis of these data identified the transcription factor KLF16 as a mediator of phenotype reversal in Atrx-deficient cells upon Sirt2 inhibition. Finally, Sirt2 inhibition impaired growth and increased senescence in ATRX-deficient GSCs *in vivo*. Our findings indicate that Sirt2 inhibition selectively targets ATRX-deficient gliomas through global chromatin remodeling, while demonstrating more broadly a viable approach to combat complex epigenetic rewiring in cancer.

**Graphical Abstract:** 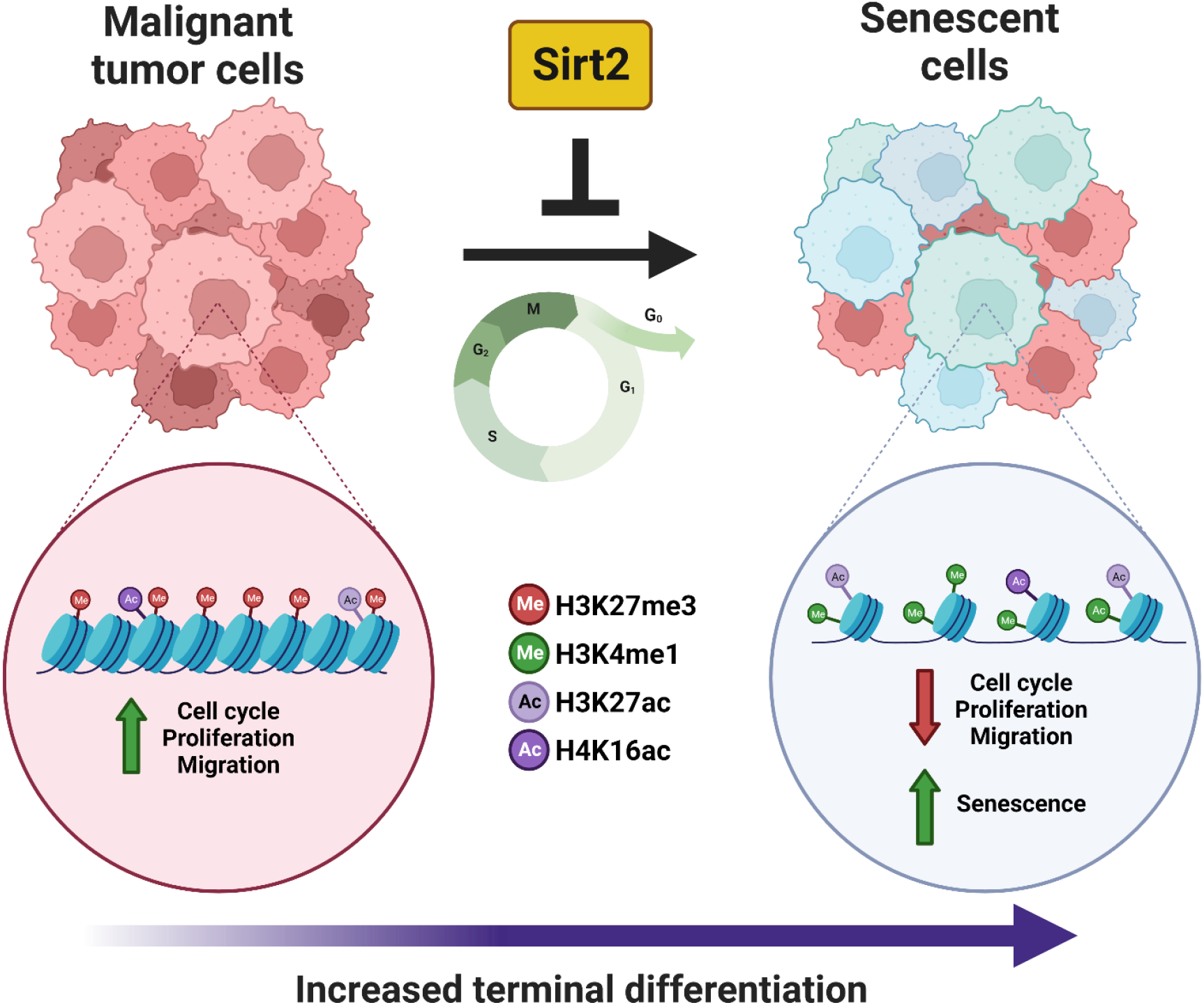

**One Sentence Summary:** Our study demonstrates that SIRT2 inhibition promotes senescence in ATRX-deficient glioma model systems through global epigenomic remodeling, impacting key downstream transcriptional profiles.

## Introduction

Malignant gliomas are the most common primary brain tumors and despite considerable molecular and clinical heterogeneity, are uniformly associated with refractivity to standard therapeutic modalities and poor overall survival(*1*). Innovative treatment strategies targeting the precise molecular mechanisms underlying glioma pathogenesis are urgently needed. Inactivating mutations in *ATRX* (α-thalassemia mental retardation X-linked), which encodes a SWI/SNF family chromatin remodeler, define major glioma subclasses also characterized by concurrent mutations in *TP53*, and in genes encoding either IDH1 or IDH2 metabolic enzymes in adults or histone H3 monomers in children(*2-5*). ATRX normally exhibits diverse functionalities and appears to play a central role in the mediation of chromatin state, gene expression, and genomic integrity in certain cell types(*6-12*). Loss-of-function mutations in *ATRX* occur at high levels across a range of neuroepithelial and mesenchymal neoplasms, where they are thought to induce alternative lengthening of telomeres (ALT), a telomerase-independent telomere maintenance mechanism(*13, 14*). ATRX deficiency also promotes replication stress and DNA damage, likely through accumulation of G-quadruplex DNA secondary structures genome-wide and modulates developmentally relevant transcriptional programs through dysregulated chromatin marks and epigenomic landscapes(*11, 12, 15, 16*). Strategies to therapeutically combat the broad physiological effects of ATRX deficiency remain unclear.

Recent work in multiple tumor variants has shown the feasibility of targeting epigenetic abnormalities to suppress oncogenic phenotypes in cancer cells, pointing to potential efficacy for analogous approaches in ATRX-deficient glioma(*17*). We previously demonstrated that ATRX deficiency induces cellular differentiation and motility phenotypes in primary murine neuroepithelial progenitor cells (mNPCs), recapitulating disease-defining characteristics of *ATRX*-mutant glioma, through the induction of genome-wide shifts in chromatin accessibility and underlying transcription(*11*). These findings suggest that sufficiently reverting the ATRX-deficient epigenomic state, along with its broad transcriptional sequelae, represents a tractable strategy for therapeutic development. To address this possibility, we coupled CMAP analysis with integrated epigenomic and transcriptional data to identify Sirt2 as a driver of ATRX-deficient oncogenic phenotypes. Targeting Sirt2 with small molecule inhibitors reverted ATRX-deficient transcriptional signatures, impairing cellular motility and promoting senescence in Atrx/ATRX-deficient mNPCs/human GSCs. Epigenomic profiling revealed that these therapeutically relevant effects were associated with shifts in H3K27ac and H4K16ac landscapes, along with underlying gene expression programs. Finally, in vivo studies demonstrated that Sirt2 inhibition impaired growth and promoted senescence in ATRX-deficient GSC xenografts. Taken together, our findings demonstrate that Sirt2-dependent epigenomic rewiring promotes ATRX-deficient oncogenic phenotypes and can be effectively targeted by small molecular inhibitors to induce senescence and terminal differentiation in *ATRX*-mutant gliomas.

## Results

### Integrated multi-omic analysis identifies Sirt2 as a targetable candidate driving ATRX-deficient phenotypes in glioma

As ATRX deficiency has been repeatedly shown to alter chromatin architecture and phenotypically relevant gene expression, we hypothesized that effective pharmacological strategies to combat ATRX-deficiency would revert ATRX-deficient transcriptional signatures. To explore this possibility, we used the Broad Institute CMAP repository to interrogate our existing RNA-seq data derived from Atrx+ and Atrx-mNPCs. CMAP compiles transcriptional signatures from cell lines subjected to either genetic or pharmacologic perturbation(*18*). Using the CLUE algorithm, we identified drug-treated gene expression signatures in CMAP demonstrating strong negative correlations with Atrx-deficent transcriptional shifts in mNPCs. Our top results were highly enriched with well-established histone de-acetylase (HDAC) inhibitors, including Vorinostat, Belinostat and Panobinostat (FIG. 1A). Moreover, we found that HDAC-associated transcriptional signatures positively correlated by gene set enrichement analysis (GSEA) with transcriptional profiles of ATRX-deficient human gliomas from the Cancer Genome Atlas (TCGA) dataset (Supplementary FIG. 1). In light of these findings, we integrated RNA-seq, ATAC-seq, and Atrx ChIP-seq data from mNPCs to determine the extent to which various HDACs were transcriptionally activated by epigenetic mechanisms in the setting of Atrx deficiency. While this analysis implicated multiple HDACs in some manner, only Sirtuin 2 (Sirt2) demonstrated positional association with an Atrx ChIP-seq peak and both increased gene expression and proximity (<2 kb) to a locus of increased chromatin accessibility—as determined by ATAC-seq—in the Atrx-deficient context (FIG. 1B-1C). Sirt2 is an NAD^+^-dependent lysine deacetylase involved in a variety of biological processes and facilitates chromatin condensation by deacetylating lysine moieties on both histone 3 (H3K27) and histone 4 (H4K16) substrates(*19-21*).

**FIG. 1.**
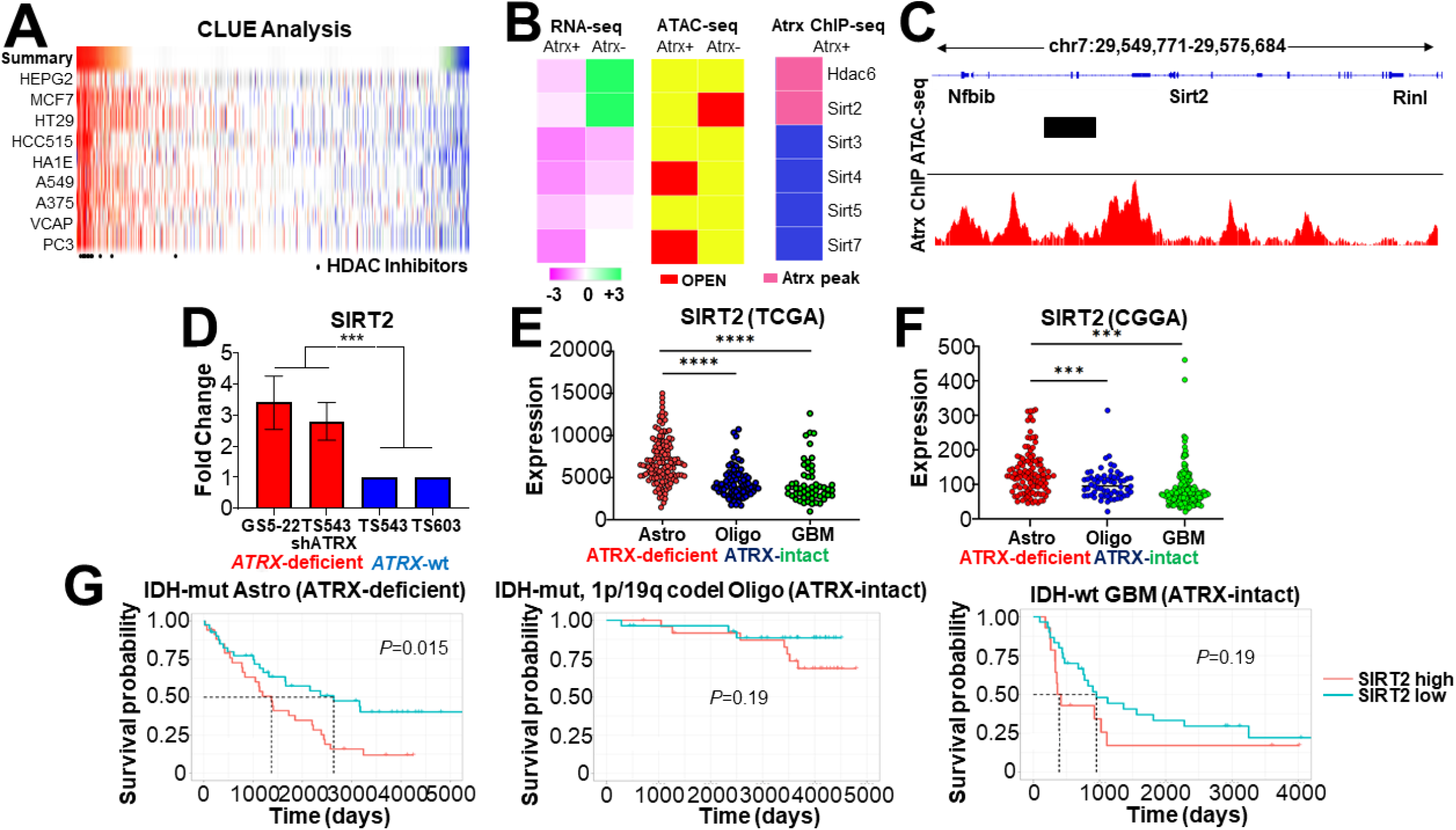

After implicating Sirt2 in ATRX-deficient gliomagenesis with our isogenic mNPCs, we sought to explore the potential clinical relevance of SIRT2 in human tumors. We first evaluated the expression of SIRT2 in a panel of patient-derived GSCs, featuring *ATRX*- mutant (GS5-22), *ATRX*-wildtype (TS543 and TS603), and *ATRX*-knockdown isogenic (TS543-shATRX) backgrounds. Using RT-PCR, we found that SIRT transcript levels were notably elevated in both *ATRX*-mutant and *ATRX*-knockdown GSCs relative to *ATRX*- wildtype isogenic and non-isogenic counterparts (FIG. 1D). We also utilized publicly available and clinically annotated gene expression data from TCGA and the Chinese Glioma Genome Atlas (CGGA). ATRX-deficient gliomas are almost exclusively found in the IDH-mutant astrocytoma subclass of adult glioma—identified by mutations in either IDH1 or IDH2 and the absence of chromosome 1p/19q codeletion (IDH-mutant astrocytomas)—where they constitute the vast majority of cases. We found that SIRT2 expression was significantly higher in this glioma subclass relative to the other major adult glioma variants (IDH-mutant and 1p/19q codeleted oligodedroglioma and IDH-wildtype glioblastoma)(FIG. 1E-1F). Moreover, when each glioma subclass was stratified by median SIRT2 transcript levels (CGGA data), high SIRT2 expression correlated with significant reduced overall survival only in ATRX-deficient gliomas with no outcome differences discerned for other disease subtypes (FIG. 1G). These data identify SIRT2 overexpression as a specific epigenetic consequence of ATRX deficiency in glioma and suggest that its therapeutic targeting may be a viable strategy for combating ATRX-deficient oncogenic phenotypes in glioma cells..

### Sirt2 inhibition attenuates Atrx -deficient transcriptional profiles impacting both cellular motility and senescence

To determine the extent to which Sirt2 activity impacts gene expression and downstream cellular phenotypes in the Atrx-deficient context, we utilized three well-characterized small molecule Sirt2 inhibitors (Sirt2i), AGK2,AK7, and Thiomyristoyl (TH). Using published IC50 values, we demonstrated increased H3 acetylation levels by western blot in Sirt2i-treated Atrx-mNPCs, an effect not seen in Atrx+ counterparts, which we had previously shown (see above) not to overexpress Sirt2 (FIG. 2A). We then used RNA-seq to assess the impact of Sirt2i on transcriptional profiles in Atrx-mNPCs, documenting significant shifts in gene expression. Importantly, treatment with each Sirt2i effectively reverted the transcriptional signature of Atrx deficiency in mNPCs by GSEA (FIG. 2B). Sirt2i-induced gene expression shifts also demonstrated associations with cytoskeletal mobilization, cell adhesion, cell motility, and senescence in Atrx-mNPCs (FIG. 2C). These latter two biological processes echo our earlier work, which identified altered cell migration and differentiation as disease-relevant phenotypes characterizing ATRX-deficient glioma.

**FIG. 2.**
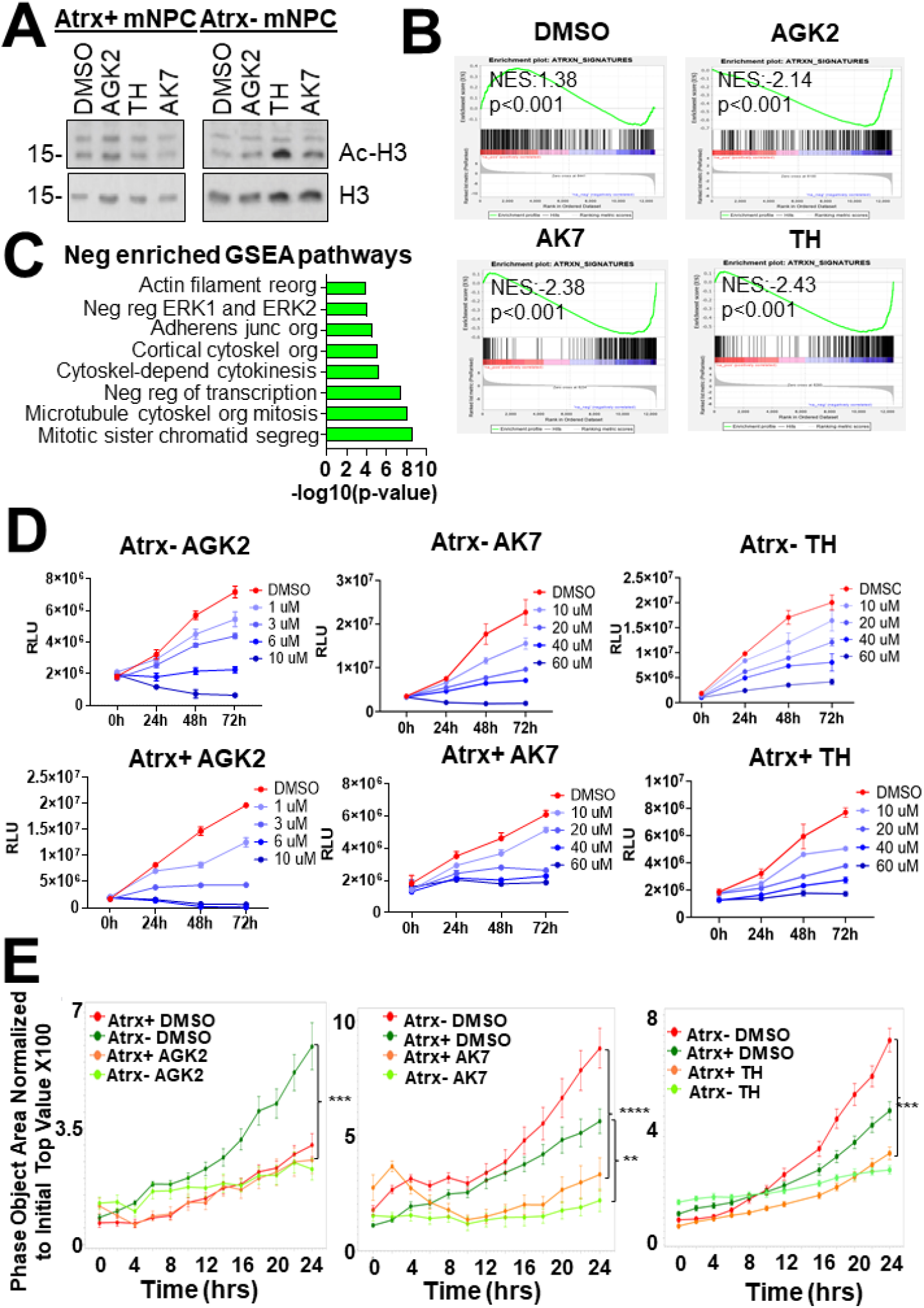

To functionally interrogate the consequences of Sirt2i-dependent transcriptional alterations, we assessed impact on cellular proliferation, migration, and senescence in murine and human experimental systems. Sirt2i decreased proliferation for both mNPCs and GSCs, regardless of Atrx/ATRX status, over a 72-hour time frame, along with mild effects on cell cycle phase distribution (FIG. 2D and Supplementary FIG. 2 and 3). These results suggest that the short-term impact of Sirt2i on glioma proliferation is modest and not ATRX-dependent. By contrast, Sirt2i exhibited much stronger influence over cell migration, profoundly impairing the motility of both Atrx-mNPCs and *ATRX*-mutant GSCs as assessed by transwell migration, with less pronounced effects on Atrx+/*ATRX*-wildtype counterparts (FIG. 2E and Supplementary FIG. 4). Sirt2i also induced β-Galactosidase, a well-established marker of senescence, to a significantly greater extent in the Atrx/ATRX-deficient context for both mNPCs and GSCs (FIG. 3A-3B). Congruent with these findings, Sirt2i treatment in mNPCs dramatically altered the levels of key senescence-associated transcripts from the SenMayo database (FIG. 3C). While the majority of SenMayo genes normally inhibit senescence and were repressed by Sirt2i, a small subset of senescence-promoting transcripts, including *Ets2* and *Sema3f*, were upregulated. Subsequent RT-PCR documented significantly increased *SEMA3F* and *ETS2* expression in AGK2-treated *ATRX*-mutant, but not *ATRX*-wildtype, GSCs (FIG. 3D). Taken together, these findings document the outsized effects of Sirt2 and Sirt2i on Atrx-deficient gene expression in glioma and associated motility and senescence phenotypes.

**FIG. 3.**
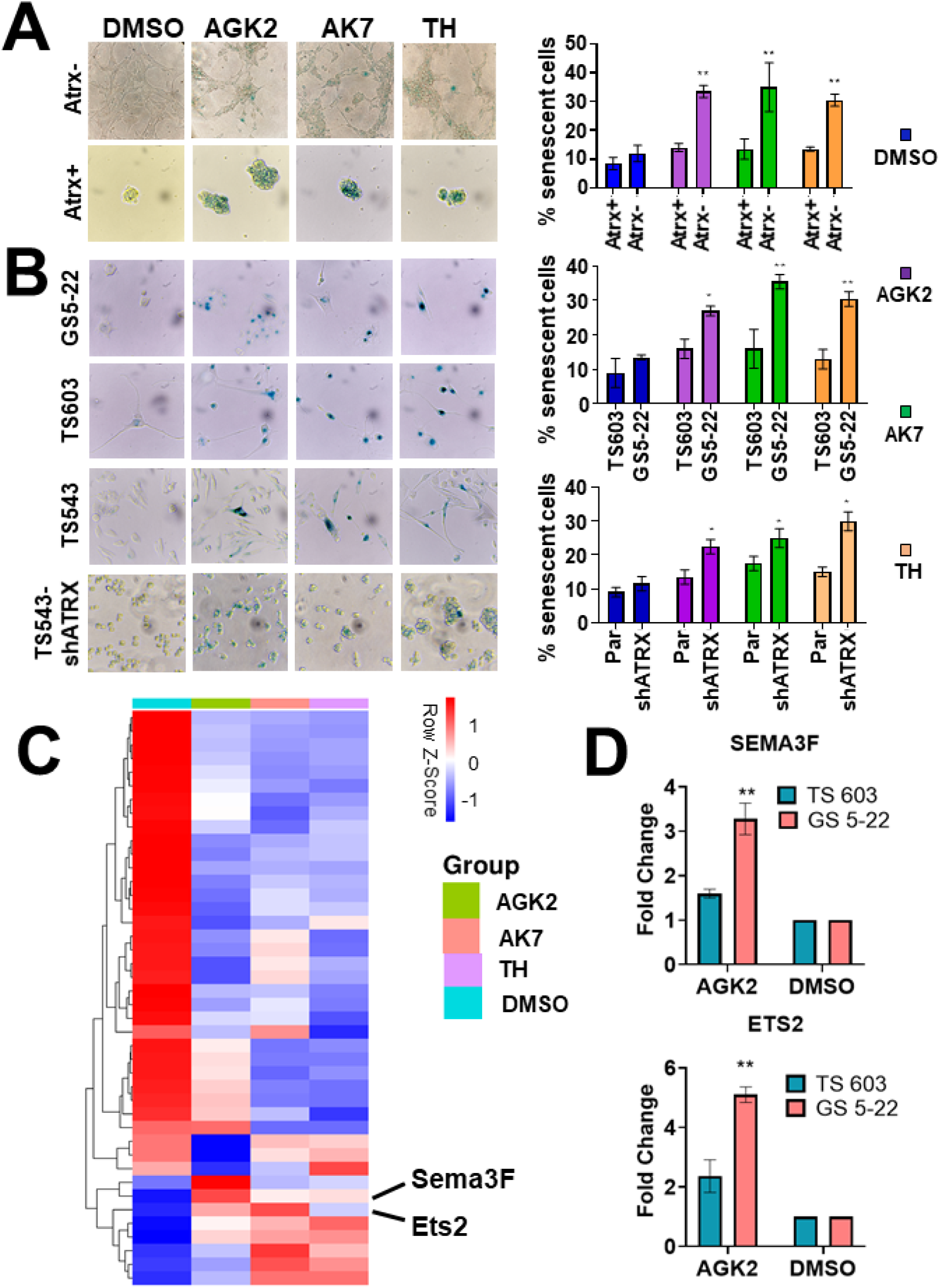

### Sirt2i induces epigenomic reprogramming in ATRX-deficient glioma models through mobilization of both H3K27ac and H4K16ac histone marks

Having documented the impact of Sirt2i on functionally relevant ATRX-deficient transcriptional shifts, we sought to delineate epigenomic mechanisms underlying these effects. As histone acetylation is typically associated with chromatin accessibility at enhancer regions, we reasoned that Sirt2i should decrease the extent of nuclear chromatin condensation. We confirmed this conjecture using single nuclei chromatin condensation parameter (CCP) analysis in mNPCs after a 48-hour treatment, observing more pronounced effects in the Atrx-context (FIG. 4A-4B). To more fully delineate alterations in enhancer landscapes genome-wide, we then performed chromatin immunoprecipitation-high throughput sequencing (ChIP-seq) on isogenic mNPCs treated with either vehicle (DMSO) or 1 µM AGK2 for 48 hours, assessing H3K27ac and H3K4me1, two marks typically associated with active enhancers, along with H3K27me3, a polycomb repression mark whose levels often demonstrate reciprocal behavior to those of H3K27ac. Sirt2i increased the size and number of H3K4me1 and H3K27ac-marked peaks with concomitant decreases in the size and number of H3K27me3-marked peaks (FIG. 4C). Once again, these effects were stronger in the Atrx-context. Applying Rank Ordering of Super-Enhancers (ROSE) to our ChIP-seq data, we found that super-enhancer (SE) calls increased more dramatically in AGK2-treated Atrx-mNPCs than in Atrx+ counterparts (3266/1158=2.82% vs 790/441= 1.79%; FIG. 4D). Pathway analysis of gene sets associated with these mobilized SEs implicated a range of relevant biological processes, including cell motility, adhesion, and death (FIG. 4E-4F). However, the subtle distinctions between these pathway correlations with regard to Atrx status, along with the absence of associated senescence signatures, raised the possibility that alternative molecular mechanisms were playing major roles in Sirt2i-induced cellular phenotypes.

**FIG. 4.**
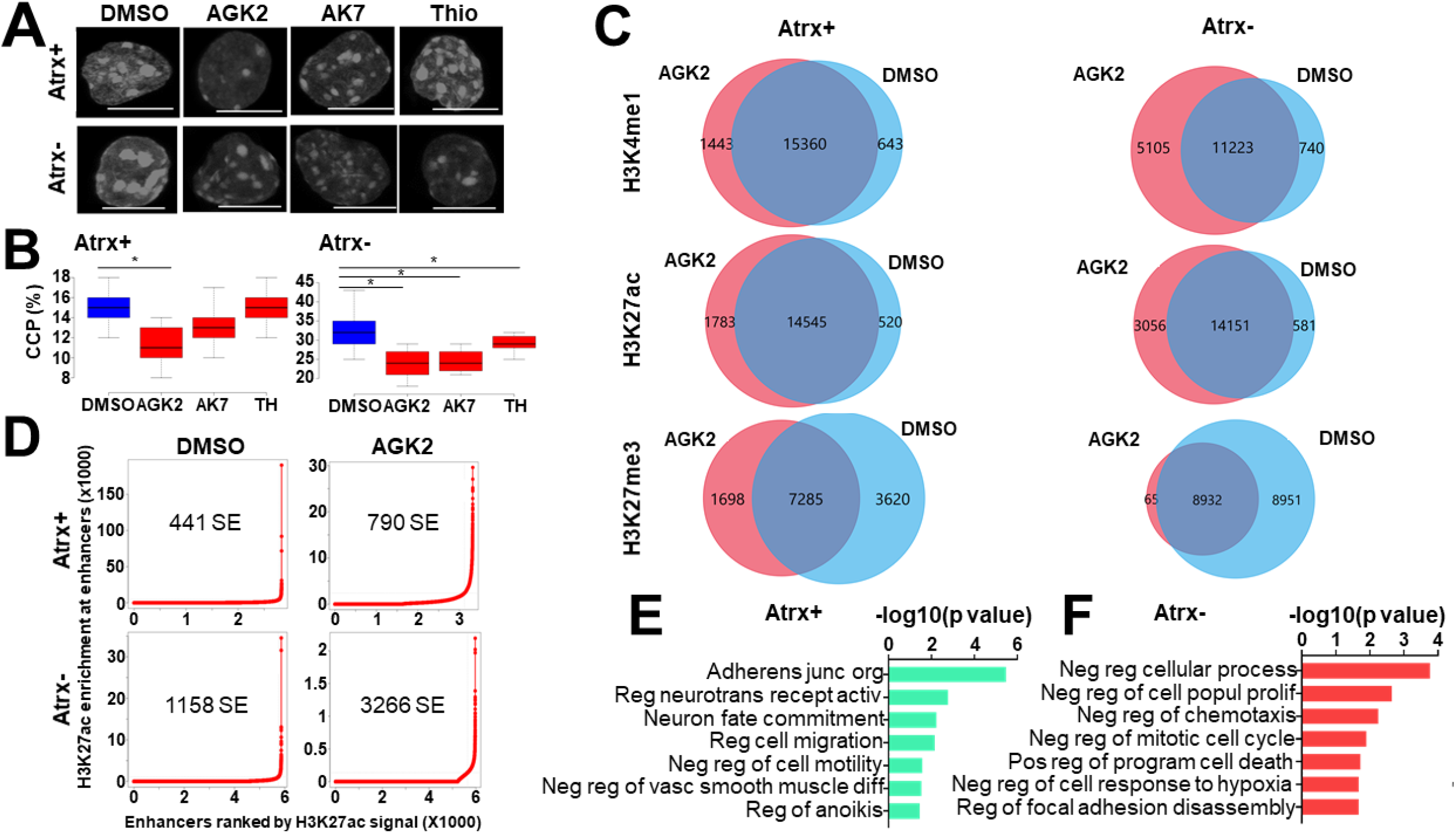

H4K16 acetylation (H4K16ac) is an enhancer-associated chromatin mark that influences both transcriptional activation and repression(*22, 23*). Intriguingly, SIRT2 has been shown to directly target H4K16ac for deacetylation during specific phases of the cell cycle(*21*). To evaluate the impact of Sirt2i on H4K16ac, we first employed immunofluorescence in mNPCs, delineating increased nuclear expression in response to drug treatment for all three inhibitors, accentuated in the Atrx-deficient setting (FIG. 5A). We then employed H4K16ac ChIP-seq in vehicle or AGK2-treated mNPCs, assessing enrichment peak number and distribution. We found that while Sirt2i had relatively modest effects in Atrx+ mNPCs, Atrx-counterparts exhibited both increased peak number and altered peak distribution (FIG. 5B). GSEA analysis of H4K16ac-marked genes arising with AGK2 treatment in Atrx-mNPCs demonstrated associations with cellular senescence, apoptosis, and adhesion not present for Atrx+ isogenics, which instead exhibited correlations with metabolic pathways (FIG. 5C).

**FIG. 5.**
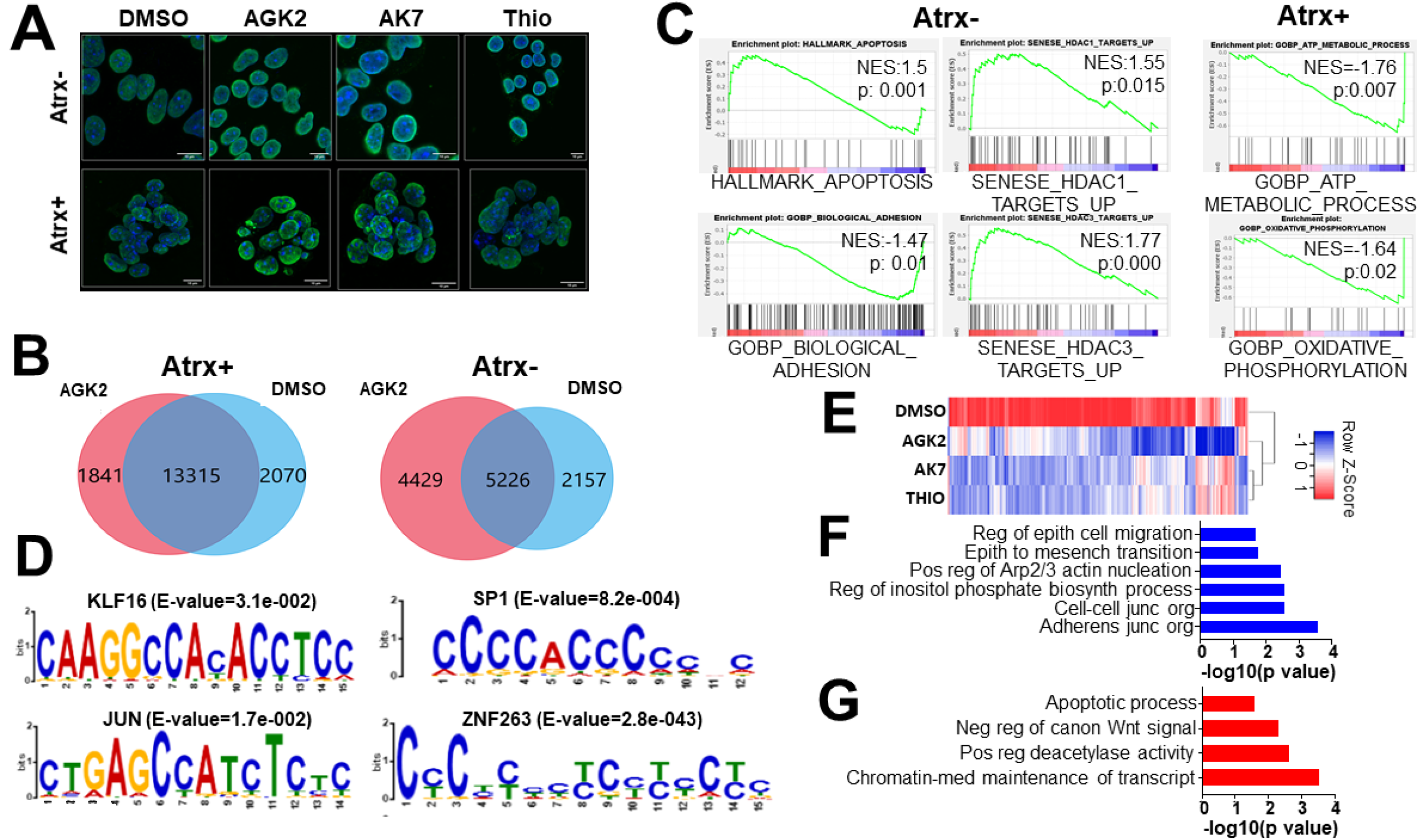

We also performed MEME analysis on AGK2-induced H4K16ac peak sites in Atrx-mNPCs to identify coregulators potentially mediating downstream transcriptional and functional consequences. This approach revealed several transcription factors (TFs) with significant motif correlations including *KLF16, JUN, SP1*, and multiple zinc finger family members (*ZNF263, ZNF384*) (FIG. 5D). Examining our RNA-seq data, we found that KLF16, a member of Krüppel-like factor (KLF) TF family and a previously reported tumor suppressor in gliomas(*24*), was specifically upregulated in Sirt2i-treated Atrx-mNPCs. Immunofluorescence in Atrx+ and Atrx-mNPCs confirmed this finding (Supplementary FIG. 5). KLF TFs bind consensus CACCC or GT boxes, induce chromatin remodeling and either activate or repress transcription based on heterodimerization partners and biological context(*25, 26*). Further emphasizing the downstream impact of KLF16 engagement by Sirt2i, we found that MEME-associated KLF16 target genes were broadly downregulated by Sirt2i in Atrx-mNPCs (FIG. 5E). GSEA analysis of these genes revealed strong negative associations with cellular motility processes and positive associations with apoptosis (FIG 5F-5G). Taken together, these findings implicate perturbed H3K27ac and H4K16ac landscapes in the pharmacological impact of Sirt2i on Atrx/ATRX-deficient glioma models. In particular, H4K16ac-based molecular mechanisms likely underlie significant effects on cell motility and senescence induced by Sirt2i in Atrx-mNPCs and *ATRX*-mutant/knockdown GSCs.

### Sirt2i impairs glioma growth and induces senescence in vivo

To evaluate the impact of Sirt2i on ATRX-deficient gliomagenesis in vivo, we generated TS543-shATRX flanks xenografts in Nu/Nu mice. Following GSC inoculation, mice were treated daily with either vehicle (DMSO) or 40 mg/kg AK7. We found that AK7 significantly slowed xenograft growth over a 15-day period and prolonged murine survival relative to vehicle treatment (FIG. 6A-6B). Moreover, immunofluorescence analysis of tumors extracted after mouse sacrifice demonstrated reduced proliferation by MIB1 (Ki67) index and increased expression of the senescence markers p16 and SEMA3F (FIG. 6C-6D). These findings demonstrate the potential of targeting SIRT2 to promote senescence and combat tumor growth in ATRX-deficient gliomas.

**FIG. 6.**
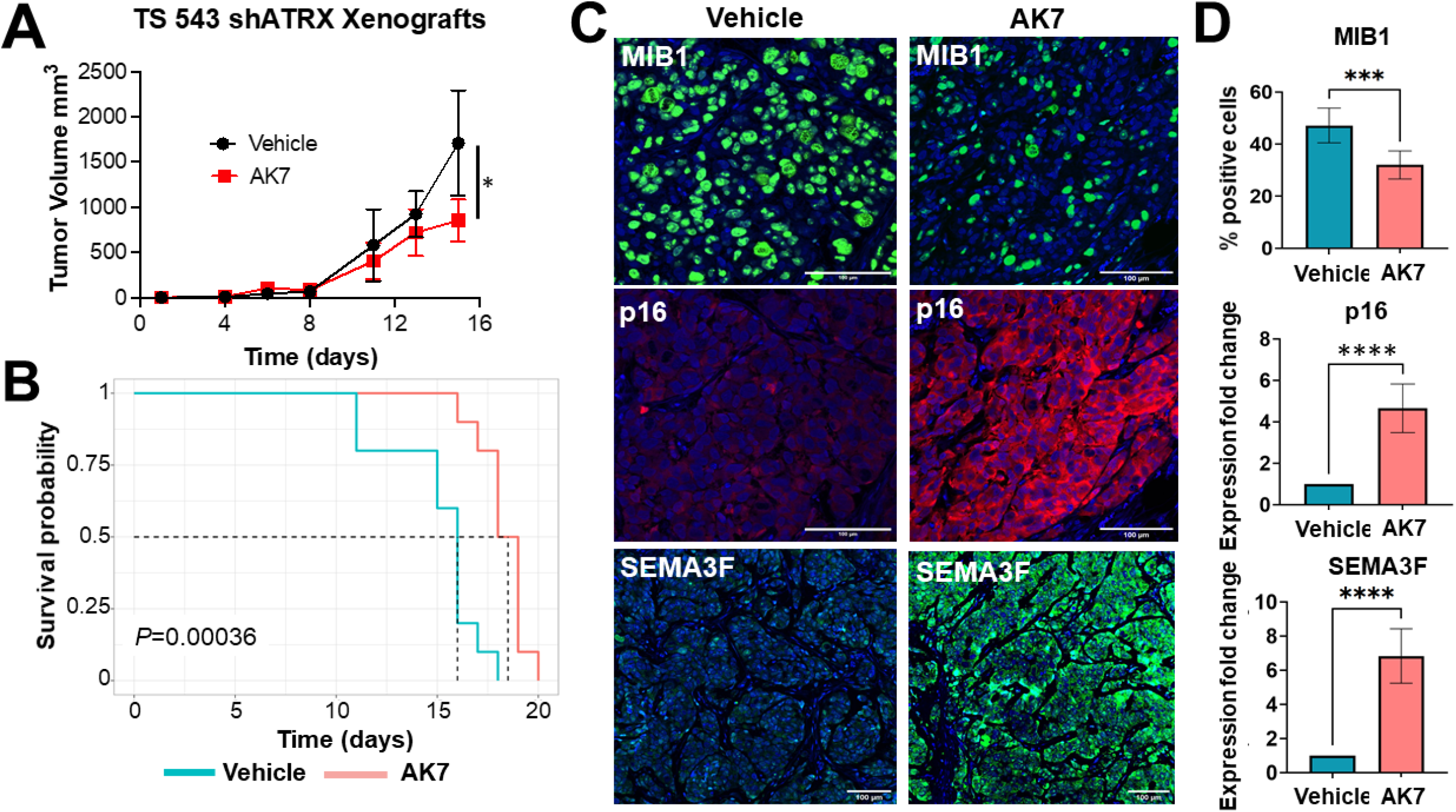

## Discussion

The discovery of epigenetic abnormalities as core features in many cancers continues to motivate efforts to characterize and target the key molecular events underlying oncogenesis in these diverse contexts. Such studies in malignant glioma are of particular need, given the dismal survival profile associated with all disease subtypes. Significant numbers of both adult and pediatric glioma feature ATRX inactivation as a defining molecular feature(*5, 27*), signaling the importance of this driver event to disease pathogenesis. However, the numerous and varied consequences of ATRX deficiency— ranging from complex chromatin remodeling to replication stress and genomic instability to telomeric dysfunction—necessarily complicate strategies for therapeutic development in affected tumors(*6, 8, 9, 12, 28, 29*). Our prior studies demonstrated that ATRX deficiency profoundly impacts chromatin accessibility genome-wide in putative glioma cells of origin, with master regular-like influence on global transcriptional profiles mediating disease-defining differentiation and motility phenotypes(*11*). In light of these findings, we reasoned that reverting the epigenomic rewiring induced by ATRX deficiency, along with downstream transcriptional sequelae, could potentially hamper its gliomagenic effects. Recent work highlights the potential of epigenetic therapies to elegantly modulate the multi-component gene sets governing complex cellular states in cancer(*30*).

Leveraging gene expression profiles for Atrx+ and Atrx-mNPC isogenics, we used the Broad Institute CMAP to identify HDAC inhibitors as promising candidates to combat ATRX-deficient epigenomic and transcriptional rewiring. Chromatin condensation by way of histone deacetylation impacts a wide range of biological processes including cell cycle, metabolism, angiogenesis, apoptosis, and differentiation(*31-33*). Small molecule HDAC inhibitors have received FDA approval for the treatment of selected hematological malignancies and have progressed through clinical trials for multiple solid tumor variants(*34*). Moreover, reported links between HDAC inhibition, terminal differentiation, and senescence induction in the glioma context were particularly intriguing in light of earlier findings from our group and others implicating ATRX deficiency in developmental disruptions within the neuroepithelial lineage(*11, 35*).

Multi-omic analyses integrating our mNPC findings with large, publicly available datasets derived from human tumors revealed the class III HDAC SIRT2 as selectively upregulated in ATRX-deficient glioma, where high expression levels selectively conferred unfavorable prognosis. Much like other HDACs, SIRT2 exhibits broad functionality across many cancer-relevant molecular networks, actively shuttling between nuclear and cytoplasmic compartments(*36, 37*). As indicated above, SIRT2 deacetylates the epigenomically active H3K27 and H4K16 histone residues, but also directly modifies a range of other targets, including α-tubulin complexes and the anaphase-promoting complex/cyclosome (APC/C)(*19-21, 38, 39*). To date, SIRT2 has been implicated in breast, non-small cell lung, hepatocellular, and colorectal carcinomas, where high expression levels are associated with aggressive clinical behavior(*37, 40*). Of note, SIRT2-mediated deacetylation of KRAS induces cell proliferation, colony formation, and tumor growth in lung and colorectal cancer models(*41*). SIRT2 also appears to regulate epithelial-to-mesenchymal transition in basal-like breast cancer by deacetylating K116 of the Slug protein(*42*). These findings speak to the multifaceted mechanisms by which SIRT2 promotes oncogenesis, as well as their intriguing cell type specificity.

Targeting SIRT2 with small molecule inhibitors has demonstrated promising results for multiple agents across several cancer models. For instance, AGK2 enhanced the anticancer effects of chemotherapeutic drugs in *TP53*-mutant colorectal cancer cells, SirReal2 induced acetylation of PEPCK1 and suppression of the RAS/ERK/JNK/MMP-9 pathway in gastric cancers, and AK7 hampered the growth of intracranial glioblastoma xenografts through increased α-tubulin acetylation(*43*). In the present study, we found that Sirt2i reverted ATRX-deficient transcriptional profiles and cell motility phenotypes, while also promoting senescence selectively in the ATRX-deficient context. These observations, initially made in mNPC isogenics, effectively translated to patient-derived GSCs. Moreover, Sirt2i also impaired ATRX-deficient GSC tumor growth in vivo, impacting both cellular proliferation and senescence. Taken together, our results demonstrate the potential clinical utility of Sirt2i in the selective targeting of ATRX-deficient epigenomic dysfunction in both adult and pediatric malignant glioma. The induction by Sirt2i of terminal differentiation in particular echoes analogous established approaches employing epigenetic therapies in myeloid malignancies and recapitulates reported effects of HDAC inhibitors in sarcoma, lung, and prostate cancer models(*44, 45*).

Impairing SIRT2 activity through pharmacological inhibition would be expected to affect a number of the protein’s downstream targets, notably impacting epigenetically influential histone modifications. Indeed, the profound shifts in global transcriptional profiles we observed with Sirt2i in Atrx-mNPCs are consistent with foundational remodeling of underlying chromatin architecture. HDAC inhibition increases chromatin acetylation levels genome-wide, directly modulating transcription factor access(*46-48*), and HDACi has been shown exert physiological effects in cancer models in large part through effects on histone acetylation(*31, 47, 49*). Consistent with this earlier literature, we found that Sirt2i lead to genome-wide increases in H3K27ac and decreases H3K27me3 in isogenic mNPCs. While these effects were more pronounced in Atrx-mNPCs, impacted genes exhibited significant correlations with similar pathways—cell adhesion and motility, apoptosis, and differentiation—in both Atrx-intact and -deficient contexts. Moreover, senescence-associated molecular networks were not implicated by GSEA analysis. Such findings were not entirely consistent with the observed phenotypic consequences of Sirt2i, particularly in Atrx-/ATRX-mNPCs and GSCs. Accordingly, we suspected that alternative epigenomic mechanisms were being engaged by Sirt2i in the Atrx-deficient context.

As mentioned earlier, SIRT2 preferentially deacetylates H4K16 during the G2/M cell cycle phase, which coincides with SIRT2 localization to cytoskeleton structures(*19-21*). Loss of H4K16ac is seen in many human cancers and aged cell states, co-occurring with site-specific DNA hypomethylation(*50-52*). H4K16ac appears to serve both enhancer and repressor functions with regard to transcription, working with the NoRC (nucleolar remodeling complex) to recruit both histone acetylases and DNA methyltransferases(*23*). We found that Sirt2i significantly and selectively enhanced H4K16ac in Atrx-mNPCs, with ChIP-seq enrichment peaks demonstrating GSEA associations with cellular senescence, apoptosis, and adhesion not present for Atrx+ counterparts. Moreover, MEME analysis implicated KLF16 as a key transcriptional regulator mediating the effects of altered H4K16ac profiles on cell motility and apoptotic gene sets. Taken together, these findings suggest that dysregulated H4K16ac promotes oncogenesis. While this unusual histone modification has yet to be extensively implicated in cancer, one very recent study demonstrated that increasing H4K16ac levels with either Ex-547, a Sirtuin 1 (SIRT1) inhibitor, or panobinostat decreased proliferative capacity in the human myeloid leukemia cells(*53*).

In summary then, we identify and characterize novel molecular mechanisms underlying the oncogenic phenotypes induced by ATRX deficiency in putative glioma cells of origin and effectively target underlying epigenomic dysfunction in murine isogenic and human GSC models, demonstrating efficacy in vitro and in vivo. Our work demonstrates a viable pathway towards precision therapeutic development for epigenetically driven cancers that leverages transcriptional signatures, experimental and in silico multi-omic analyses, and repurposed pharmaceuticals. We expect that analogous approaches could be effectively implicated in other deadly tumors.

### Methods

#### CLUE data analysis

CLUE (https://clue.io/) is a cloud-based hub designed to identify candidate drug compound by way of transcriptional signatures(*18*). Genes differentially expressed between Atrx- and Atrx+ mNPCs, were mapped, ranked, and filtered for high fold changes (<1.5 fold change) and low p-values (-<0.01) and matched to human orthologs using the DAVID conversion tool (https://david.ncifcrf.gov/conversion.jsp). These signatures were then correlated with transcriptional profiles generated by perturbagen exposure in nine human cancer cell lines (HEPG2, MCF7, HT29, HCC515, HA1E, A459, A375, VCAP and PC3), using a comprehensive ranking score system.

### Cell lines

All cell lines used in this study were tested for mycoplasma contamination in every three months and used a minimal passage number. mNPCs (Atrx+ and Atrx-) were generated as described previously(*11*) and cultured in NeuroCult Basal Medium containing NeuroCult Proliferation Supplement, 20 ng/ml EGF, 10 ng/ml basic FGF, 2 μg/ml heparin (Stemcell Technologies). GS 522, TS 603, TS 543 and TS 543-shATRX have been described previously(*11, 12*) and were cultured in DMEM/F12 media with 20 ng/ml EGF, 10 ng/ml basic FGF, 2 μg/ml heparin (Stemcell Technologies). The status of ATRX, *TP53* and *IDH1* (R132H) were routinely assessed in all lines either western blot or Sanger sequencing.

### Quantitative reverse transcriptase PCR (RT-qPCR)

Total RNA was extracted using Qiagen RNeasy Plus, followed by DNAse treatment, as per manufacturer’s instructions. 1 γg of total RNA from each sample was then converted to cDNA using First Strand cDNA Synthesis Kit (#K1612, Thermo), followed by qPCR using Power SYBR Green PCR Master Mix (Applied Biosystems). Data was analyzed by the ΔΔCt method using GAPDH as a housekeeping gene. Primers for RT-qPCR are listed in Supplementary Table 1.

### Sirt2 inhibitors and cell viability assays

AGK2, AK7 and Thiomyristoyl were obtained from SelleckChem. Assays incorporated 5000 cells/well with serial concentrations for AGK2 (1uM-10uM), AK7 and Thiomyristoyl (10uM-60uM) incubated for 48 hours in 96-well plates. Viability was then assessed with the CellTiter-Glo Luminescent assays (Promega) according to manufacturer-recommended procedures and quantified using a Tecan INFINITE M1000 PRO luminometer.

### Cell migration assays

mNPCs and human GSC lines were seeded on laminin (20 μg/ml) coated 96 well dishes. Cells were grown into a dense monolayer and then scratched with the Wound Maker (Essen BioScience), following manufacturer instructions. Sirt2i (AGK2: 1uM, AK7 and Thiomyristoyl: 10uM) and DMSO were added with appropriate media and wound closure was monitored using an IncuCyte Zoom System (Essen BioSciences) for 48 hours. Images were analyzed with Incucyte Scratch Wound Cell Migration Software (Essen BioSciences).

### RNA-seq and data analysis

Atrx-mNPCs were treated with Sirt2 inhibitors (AGK2: 1uM, AK7 and Thiomyristoyl: 10uM) or DMSO vehicle for 48 hours, followed by total RNA extraction using Qiagen RNeasy Plus kit (three biological replicates), according to manufacturer’s instructions. Subsequent processing and sequencing occurred at the MD Anderson Advanced Technology and Genomics Core, (ATGC). Following Agilent BioAnalyzer quality assessment, libraries were generated using Truseq library preparation kits (Illumina) and samples run on the HiSeq4000 platform in a 76bp pair-end sequencing format. The raw fastq files were subjected to FASTQC analysis for quality control analysis, followed by alignment to mouse mm9 genome using RNASTAR (version 2.7.8a). Raw transcript counts were generated using HTseq-count tool (version 0.9.1), followed by Principal Component Analysis (PCA). Differentially expressed genes (DEGs) were calculated using DESeq2 (version 2.11.40.7) and were later subjected to Gene Set Enrichment Analysis (GSEA) analysis using a desktop version of the analysis tool.

### Antibodies

All commercial antibodies utilized for ChIP-seq, western blotting and immunofluorescence in the study are listed in Supplementary Table 2.

### ChIP-seq analysis

Atrx-mNPCs were treated with either AGK2 (1uM) or DMSO for 48 hours, fixed with 37% formaldehyde, and quenched with 125 mM glycine. Samples were then sonicated using a Bioruptor Plus (Diagenode) and immunoprecipitated using antibodies to H3K4me1, H3K27ac, H3K27me3, and H4K16ac. DNA was reverse-crosslinked for 4 hours, followed by solid-phase reversible immobilization (SPRI) beads cleanup and eluted in Tris-EDTA buffer. Eluted DNA was assessed for quality, followed by library preparation and run on a Novaseq6000 using a 50bp single-end read format at the ATGC. Generated ChIP-seq data underwent quality-control measures using the FASTQC tool and aligned to the mm9 reference genome with Bowtie2 (version 2.4.5). Only unique mapped reads were retained. MACS2 callpeak (version 2.2.7.1) was used for calling peaks from BAM files with default parameters. BED files were further subjected to gene annotation using GREAT (version 4) analysis with Basal plus extension parameters (Proximal: 5kb upstream, 1kb downstream, plus distal upto 1000kb). For MEME analysis, ChIP-MEME was utilized using default parameters. Peak annotation and visualization were performed using the PAVIS annotation tool. ChIP-seq heatmaps were generated using seqMINER. For Super-enhancer calling, ranking of the super-enhancers (ROSE) algorithm was used on H3K27ac ChIP-seq data, followed by pathway prediction analysis using Enrichr.

### Immunofluorescence analysis

For immunostaining, mNPCs were grown on chamber slides treated with Sirt2i (AGK2: 1uM, AK7 and Thiomyristoyl: 10uM) for 48 hours and were fixed using 4% paraformaldehyde for 10 mins. Cells were permeabilized using 0.5% Tween-20, 0.2% Triton X-100 in PBS for 10 min and blocked for 1 hour 5% BSA solution, followed by overnight incubation with primary antibody. Slides were washed 3 times with PBST solution and incubated with Alexa Fluor 488/594 secondary antibodies (Invitrogen) at room temperature for 1 hour. Slides were washed thrice and mounted with coverslips using DAPI counterstain (#H-1200-10, Vector laboratories). Imaging was done using an Olympus confocal microscope (FV-1000), Z-stack images were captured using a 60× objective with a pixel size of 0.1-µm and 0.5-µm depth. Images were processed using Fiji software.

### Chromatin Condensation Parameter (CCP) analysis

Measurement for chromatin condensation was performed using single nuclei DAPI stained images for mNPCs treated with Sirt2i (AGK2: 1uM, AK7 and Thiomyristoyl: 10uM) for 48 hours. CCP was developed in MATLAB and utilized the Sobel edge detection algorithm for analysis. Z-stack images were thresholded and processed using Fiji software followed by CCP analysis.

### Protein isolation and Western blot

mNPCs were treated with Sirt2i (AGK2: 1uM, AK7 and Thiomyristoyl: 10uM) for 48 hours, lysed using RIPA buffer (150 mM NaCl; 50 mM Tris pH 8.0; 1.0% IGEPAL CA-630; 0.5% sodium deoxycholate; 0.1% SDS; Sigma-Aldrich) supplemented with protease inhibitors (cOmplete mini, Roche Diagnostics; 1 mM PMSF; 10 mM NaF; 2.5 mM Na3VO4) and centrifuged at 10,000 × g at 4 °C for 30 min. The resulting supernatants were quantified using BCA protein assays (#23225, Thermo) and immunoblotted with standard procedures.

### Senescence β-Galactosidase Staining

Sirt2i (AGK2: 1uM, AK7 and Thiomyristoyl: 10uM) or vehicle-treated mNPCs and human GSCs were treated for 48 hours. The cells were fixed using 4% paraformaldehyde for 10 mins using the Senescence β- Galactosidase Staining Kit and stained for β-galactosidase overnight following manufactures instructions (#9860, CST). Cells were then washed with PBS twice and β- galactosidase-positive constituents counted at low magnification (20x) and images were taken using Nikon ECLIPSE Ts2 inverted microscope and processed using Fiji software.

### Mouse xenograft experiments

All animal procedures described herein were approved by Institutional Animal Care and Use Committee (IACUC) of M.D. Anderson Cancer Center (protocol # 00001597-RN01). TS543-shATRX GSCs were grown in cell culture dishes, dissociated with Accutase (#07920, Stemcell), resuspended in DMEM/F12 media mixed 1:1 with Matrigel Growth Factor Reduced (GFR) Basement Membrane Matrix (#354230, Corning), and injected into nude mice flanks in a 50 μl aliquot containing 2×10^6^ cells. Mice were randomly segregated into two groups: vehicle or AK-7 (40mg/kg), each administered via intraperitoneal injection, daily until health-defined end points. Tumor size was measured with calipers. For growth curve analysis and survival studies, mice in the study cohorts were sacrificed when tumors reached 2000 mm^3^ in total volume. The tumors were excised from sacrificed mice and subjected to routine histopathological processing

### Immunohistochemistry

Five-micron FFPE sections were deparaffinized and subjected to antigen retrieval with citrate buffer (10mM). Sections were then blocked with 2% goat serum for 1 hour, followed by overnight incubation with primary antibody at 4°C. Sections were washed with 0.01% PBST and incubated with ImmPRESS-HRP Goat Anti-Rabbit IgG secondary antibody (Vector Laboratories) and incubated for 1 hour, and developed using Alexa Fluor 488/594 Tyramide SuperBoost Kit. Slides were washed thrice with 0.01% PBST and mounted with coverslips using DAPI counterstain (#H-1200-10, Vector laboratories) and imaged using an Olympus confocal microscope (FV-1000). Images were processed using Fiji software.

### TCGA and CGGA gene expression data

Level-3 gene expression data was obtained from TCGA along with the corresponding sample annotation file (https://www.cancer.gov/about-nci/organization/ccg/research/structural-genomics/tcga).

For CGGA, count data and clinical information were downloaded from the access portal (http://www.cgga.org.cn/download.jsp).

### Statistics

All statistical analyses were performed with either GraphPad Prism 8.0, R (v3.5.0) or Microsoft Excel. All results, unless otherwise stated, represent at least three independent experiments and are plotted as mean ± SEM. All data were statistically analyzed using unpaired or paired, two-tailed t-tests. For transcriptional and epigenome data, the hypergeometric distribution was utilized and followed default statistical parameters. P values are represented using * for P < 0.05, ** for P < 0.01, *** for P < 0.001, and **** for P < 0.0001.

## Supplementary Figures

**Supplementary FIG. 1:**
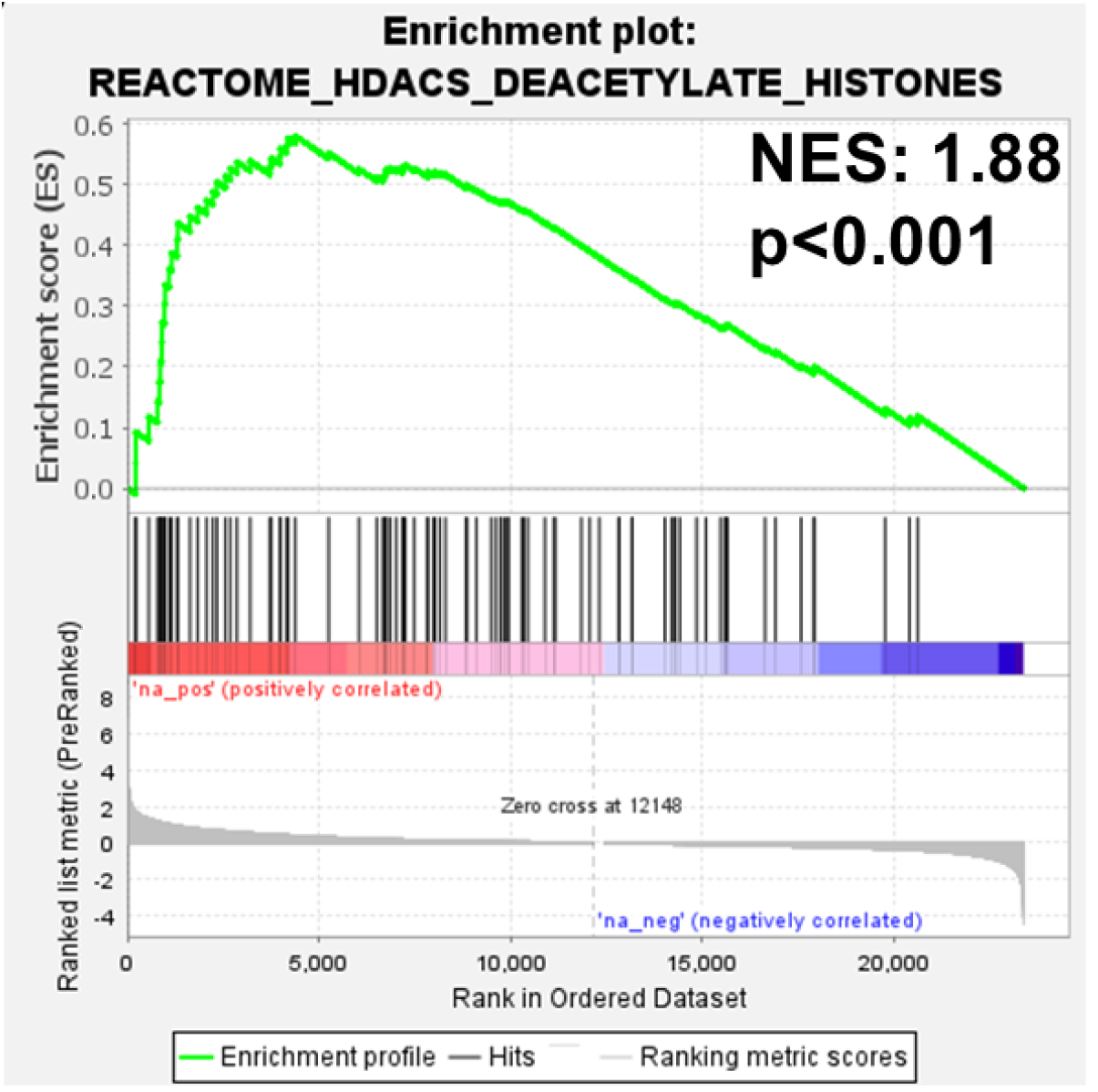
GSEA plot demonstrating HDAC-associated transcriptional signatures enriched in ATRX-deficient human gliomas from TCGA data.

**Supplementary FIG. 2:**
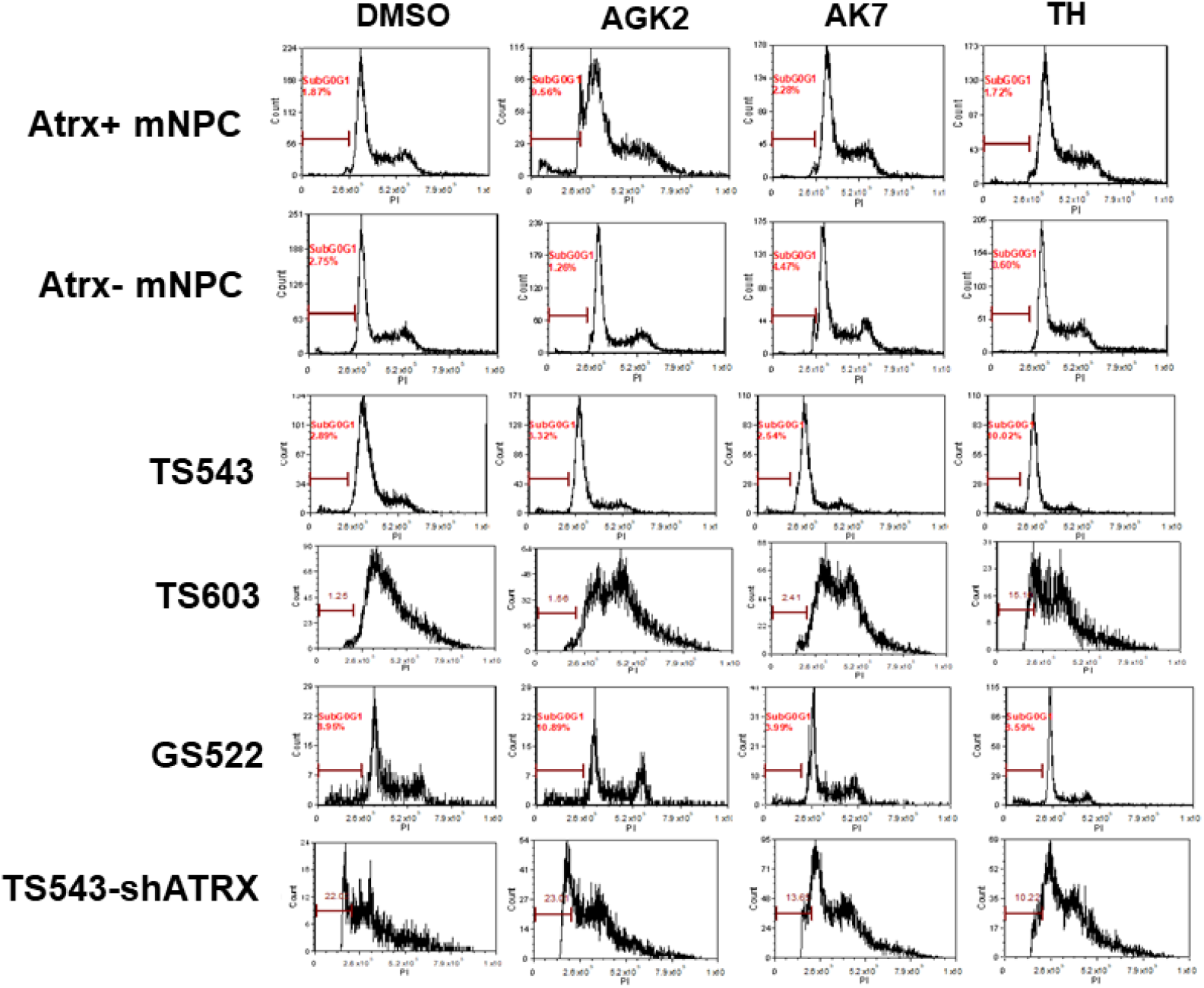
Cell cycle analysis for mNPCs and GSCs treated with either Sirt2i or DMSO (vehicle) for 48-hours showing modest effects cell cycle phase distribution.

**Supplementary FIG. 3:**
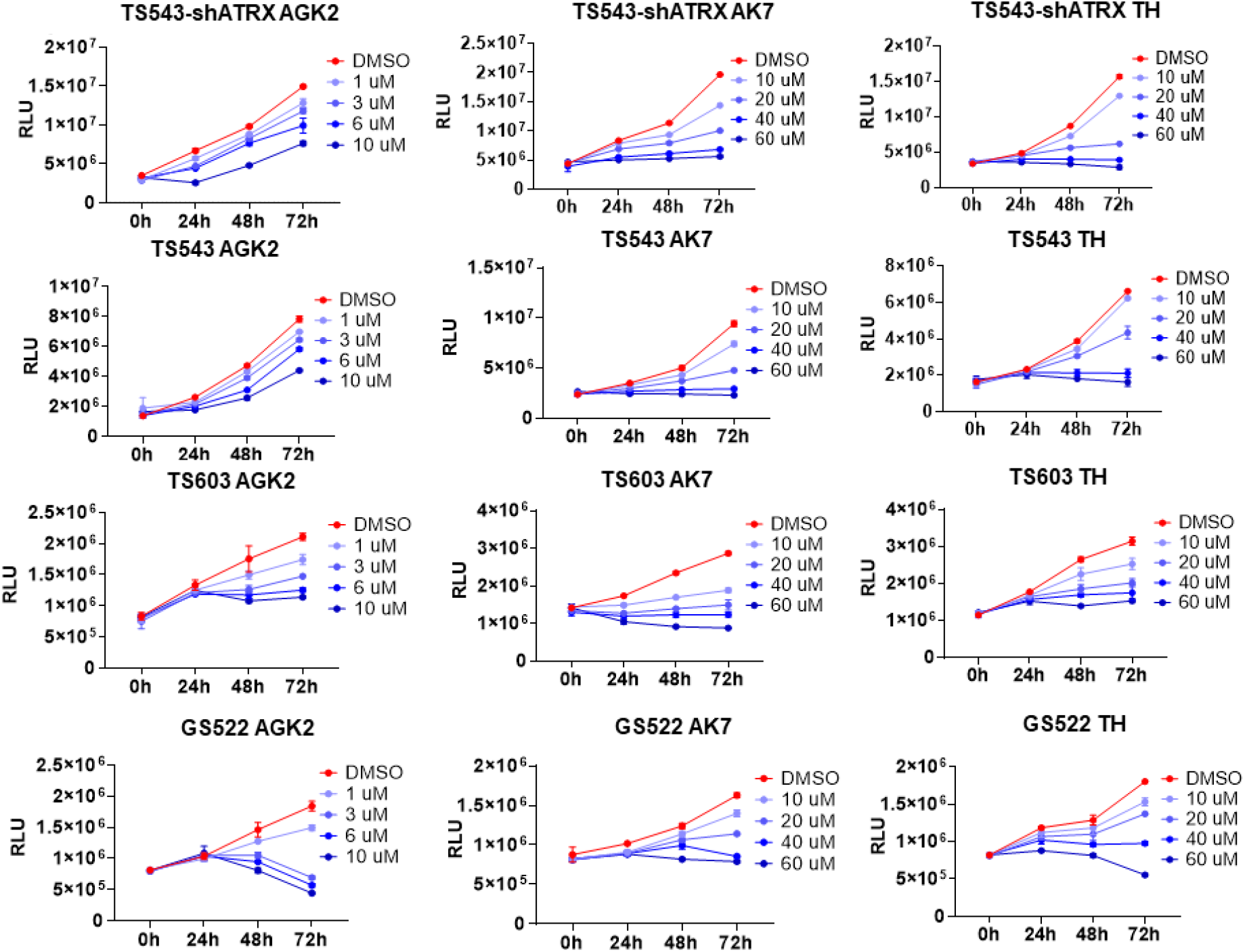
Cell proliferation analysis for GSCs treated with either Sirt2i or DMSO (vehicle) for 24, 48 and 72-hours.

**Supplementary FIG. 4:**
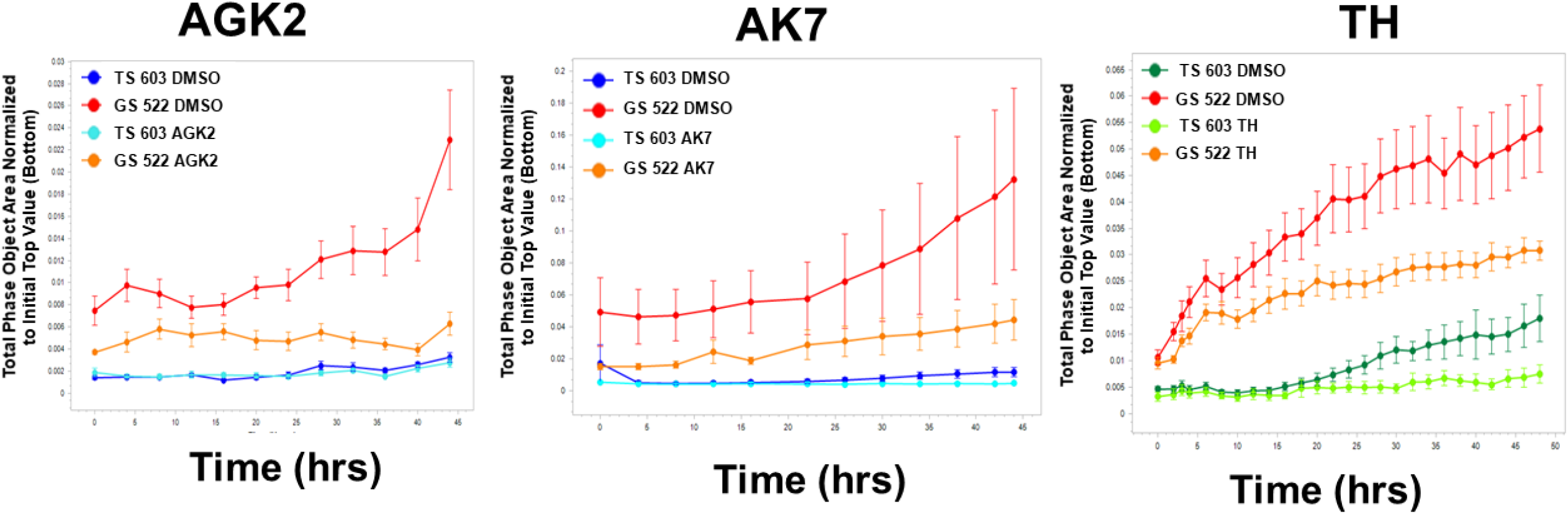
Cell migration analysis for GSCs treated with either Sirt2i or DMSO (vehicle) for 48 hours.

**Supplementary FIG. 5:**
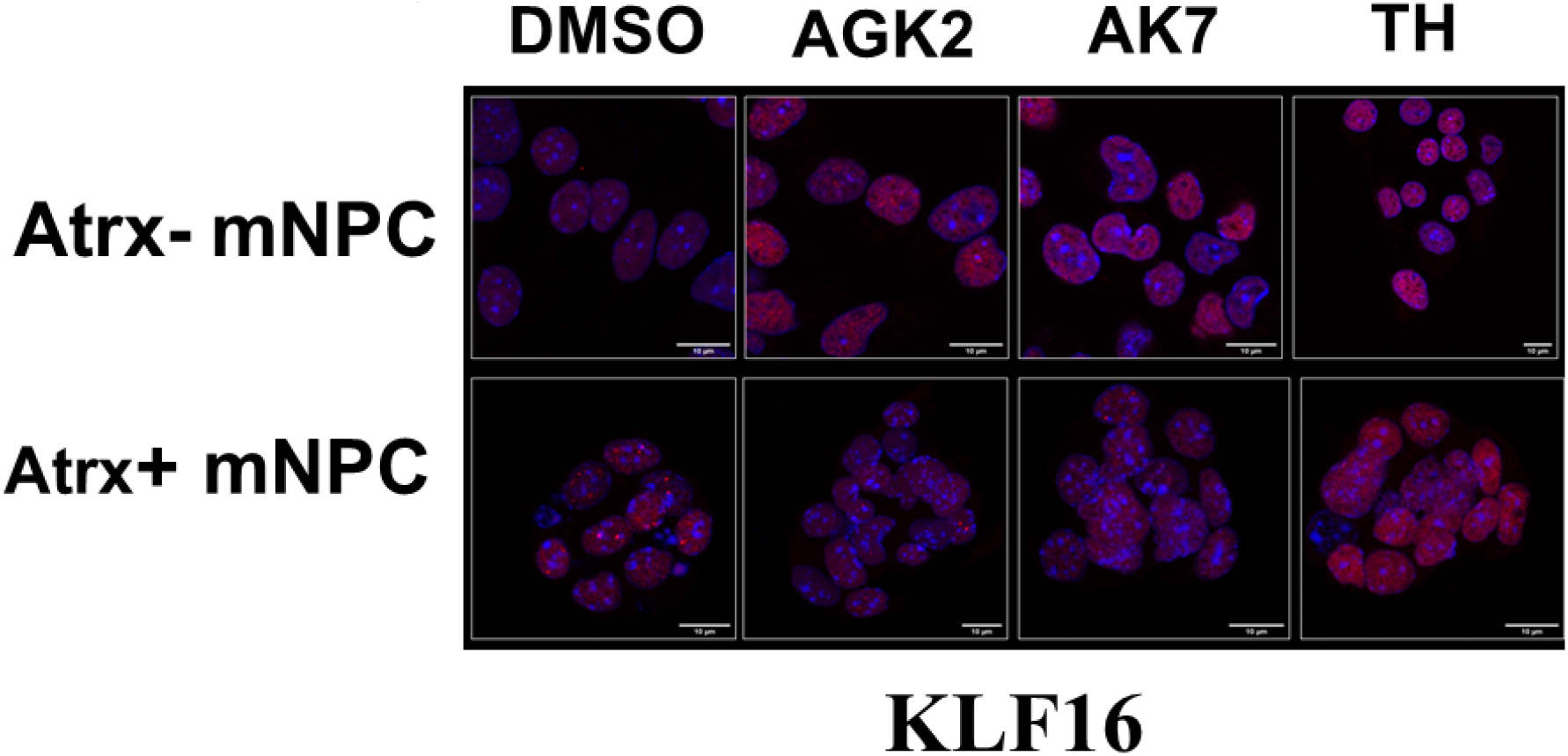
Immunofluorescence analysis for KLF16 in mNPCs treated with either Sirt2i or DMSO (vehicle) for 48 hours demonstrating increased expression exclusively in Atrx-deficient isogenics.

## Supplementary Table

**Supplementary Table.**
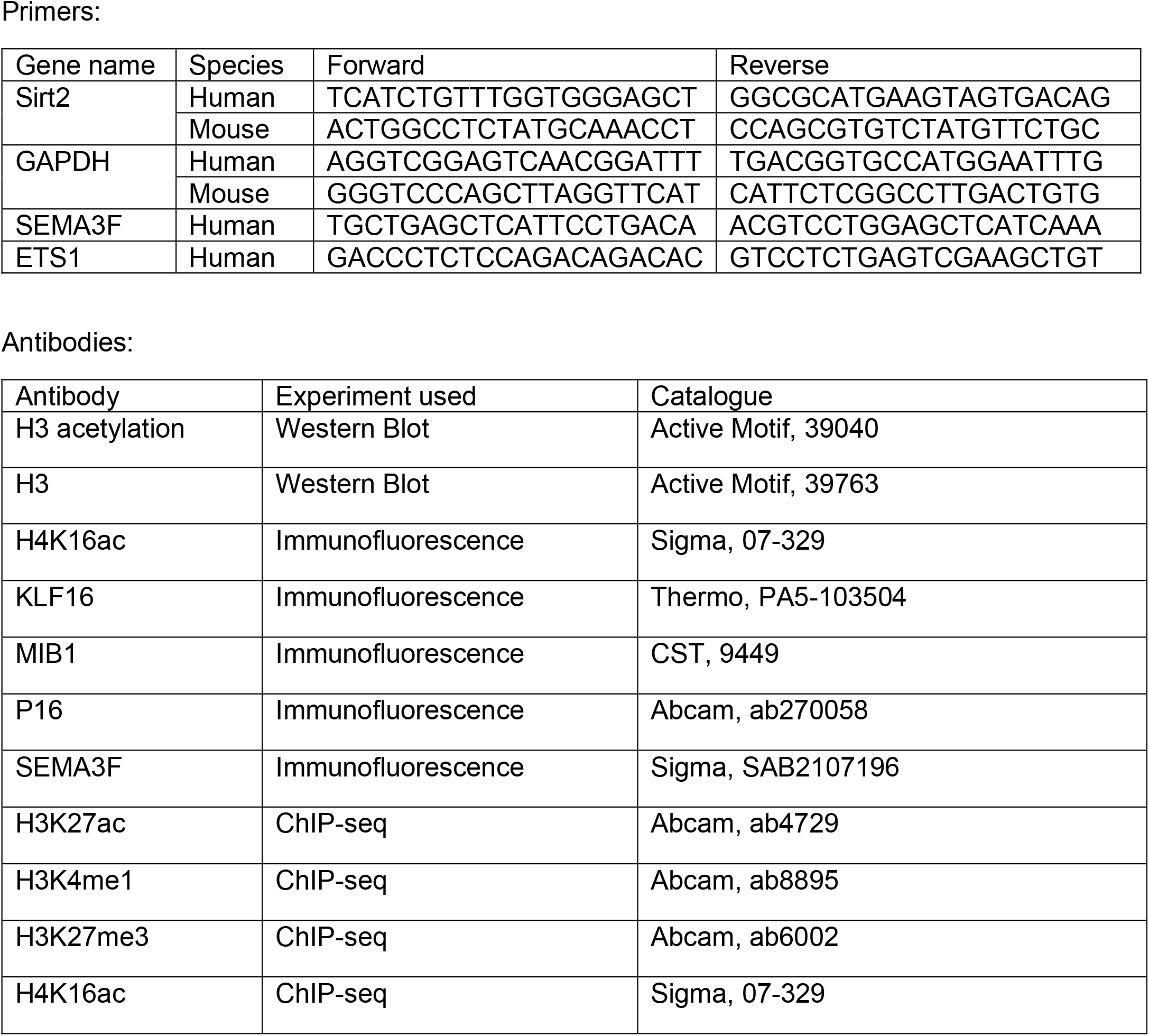
List of primers and antibodies utilized in this study.

## Acknowledgments

We would like to graciously thank Drs. Frederick Lang and Erik Sulman for providing access to GSC lines. We would also like to thank Jacob Mattia for assistance with CCP experiments. Finally, we would like to acknowledge the MD Anderson Genomics Core Facility (GCF).

## Author contributions

Conceptualization: PBM, CD, JTH; Methodology: PBM, CD, SD, AR, JTH ;Investigation: PBM, CD, SD, AR, JTH; Funding acquisition: JTH; Project administration: PBM, JTH Supervision: PBM, JTH ;Writing – original draft: PBM, JTH ; Writing – review & editing: PBM, JTH

## Data and materials availability

All transcriptional and epigenomic data have been deposited in the Gene Expression Omnibus (GEO) repository under accession number ----.

